# Cohesiveness in microbial community coalescence

**DOI:** 10.1101/282723

**Authors:** Nanxi Lu, Alicia Sanchez-Gorostiaga, Mikhail Tikhonov, Alvaro Sanchez

## Abstract

Microbial invasions exhibit many unique properties; notably, entire microbial communities often invade one another, a phenomenon known as community coalescence. In spite of the potential importance of this process for the dynamics and stability of microbiome assembly, our understanding of it is still very limited. Recent theoretical and empirical work has proposed that large microbial communities may exhibit an emergent cohesiveness, as a result of collective consumer-resource interactions and metabolic feedbacks between microbial growth and the environment. A fundamental prediction of this proposal is the presence of ecological co-selection during community coalescence, where the invasion success of a given taxon is determined by its community members. To establish the generality of this prediction in experimental microbiomes, we have performed over one hundred invasion and coalescence experiments with environmental communities of different origins that had spontaneously and stably assembled in two different synthetic aerobic environments. We show that the dominant species of the coalesced communities can both recruit their community members (top-down co-selection) and be recruited by them (bottom-up co-selection) into the coalesced communities. Our results provide direct evidence that collective invasions generically produce ecological co-selection of interacting species, emphasizing the importance of community-level interactions during microbial community assembly.

## Introduction

Ecologists have long contemplated the idea that interactions between two or more co-invading species can produce correlated invasional outcomes ^1–6^. For instance, the hypothesis known as “invasional meltdown” proposes that facilitative interactions between co-invading species can amplify their negative effects on the resident community, enhancing their own invasive success and facilitating future invasions ^7–9^. These effects are believed to be particularly strong during invasions by ecosystem engineers, those species that actively alter the environment and which, by doing so, may destroy existing niches in the resident ecosystem ^1,8^.

Recent research has highlighted that co-invasions are very common in the microbial world^10–14^. The human digestive system, for instance, can get invaded several times a day by the microbial communities that reside on the ingested food or drink. This phenomenon, where entire microbiomes invade one another, has been termed microbial community coalescence, and is reminiscent of the sudden contact between previously isolated ecosystems when a geographical barrier that separated them is suddenly removed ^3,10^. In spite of its clear potential importance, the role of community coalescence in microbiome assembly is only beginning to be addressed ^10^, and little is known about its potential implications and mechanisms.

Early mathematical models of community-community invasions ^3,15^, as well as more recent theoretical work that includes environmental construction (e.g. consumer-resource models) ^13,14^, both suggest that high-order invasion effects can be expected during community coalescence. Communities that are not randomly formed and have a previous history of coexistence may exhibit an emergent “cohesiveness”, which produces correlated invasional outcomes amongst species from the same community ^4^. For instance, the metabolically more efficient community might be expected to collectively dominate after community coalescence, since it will overwhelmingly affect the environment and lead to conditions which its members can tolerate, but the members of the other community generally may not ^14^. Results from community coalescence experiments in single-batch anaerobic fermenters have been found to be consistent with this prediction ^12^, and with the existence of correlated invasional outcomes. This latter phenomenon has been termed “ecological co-selection” ^11,12^, to reflect that ecological partners in the invading community recruit one another into the final coalesced community.

The mechanics of ecological co-selection during community coalescence are still poorly understood. Is it governed largely by just a few key species, which recruit everyone else, or by collective interactions among all, including the rarer members of the community? Discerning between these two possibilities is a question of fundamental importance. While natural microbial communities are highly diverse, they are typically also very uneven, and the role played by the rare species has long been the subject of debate ^16^. Due to their large population sizes, it is reasonable to expect the dominant species to have a proportionally oversized influence on shaping the environment in their communities. For instance, laboratory cultures often contain strains that feed off the metabolic secretions of the dominant species ^17^. In such a scenario, the fate of the dependent sub-dominant species might be tied to the success of their dominant taxon, which provides the required nutrients. In an alternative scenario, the most abundant taxon in a community might owe its dominance, at least in part, to cross-feeding or other forms of facilitation from the rarer members of the population. Therefore, in the context of coalescence experiments, one could imagine scenarios of either “top-down” cohesiveness, where the dominant taxa influence the invasion success of the the rarer members of the community; and “bottom-up” cohesiveness, where the rarer or subdominant members of a community influence the invasion ability of their dominant member. Either of these two forms of ecological co-selection could in principle be positive (recruitment) or negative (antagonistic) (See Fig. 1E). Concrete ecological and molecular mechanisms could be proposed for either of the four scenarios, but which of them are typically realized in natural communities? Addressing this question has been experimentally challenging in the past ^11,12^.

**Figure 1.**
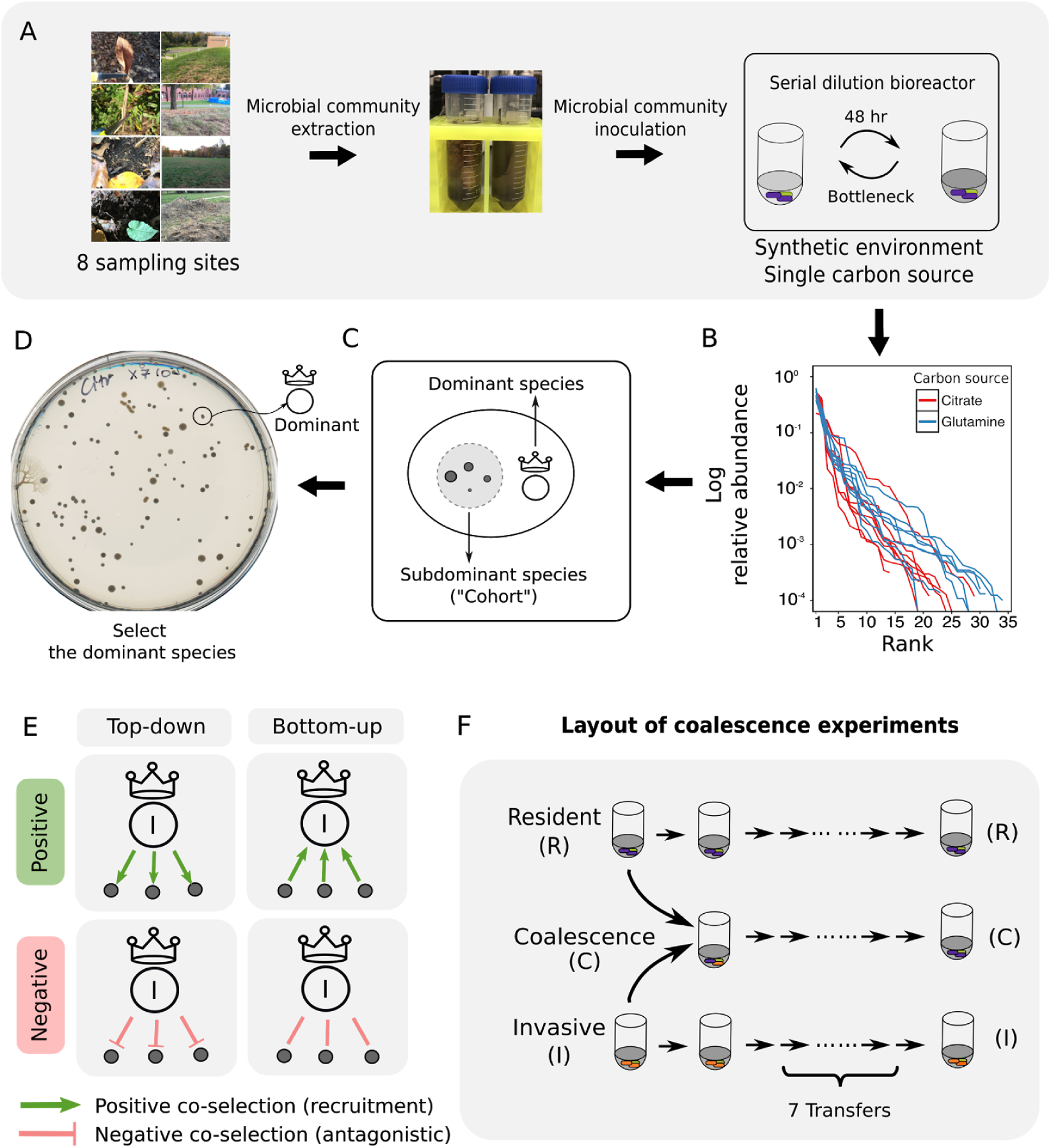
Outline of experimental strategy. (A) Microbial communities were isolated from eight different locations and stabilized in serial batch culture bioreactors as described elsewhere ^18^. (B) Rank-Frequency distributions for all eight communities reared in either citrate or glutamine as the only carbon source, sequenced at a depth of 10^4^ reads. Community composition is skewed and long-tailed. (C-D) The dominant (i.e. most abundant) species of each community were isolated on minimal media agar plates. The “cohort” is the collection of all sub-dominant species in a community. (E) Our hypothesis: ecological co-selection can be “top-down”, where the dominant co-selects the cohort; or “bottom-up”, where the cohort co-selects the dominant. Both forms of co-selection can be positive (recruitment), or negative (antagonistic). (F) Depiction of our coalescence experiments.

In previous work, we have shown that large numbers of soil and plant microbiomes can be cultured *ex situ* in synthetic minimal environments with a single supplied limiting resource under serial growth-dilution cycles (Fig. 1A)^18^. Under these conditions, environmental microbiomes spontaneously re-assemble into complex multi-species communities that are stabilized by dense cross-feeding facilitation networks ^18^. These aerobically growing communities are easy to manipulate; grow in high throughput; rapidly stabilize under serial daily transfers; are largely made up by culturable members; and strongly engineer their own environment (e.g. upwards from 40% of their total biomass can be produced from secreted metabolic byproducts) ^18^. Because of this we reasoned that they are good test-cases to investigate ecological co-selection during community coalescence.

In addition, just like natural assemblies, our communities are typically long-tailed and uneven (Fig. 1B, Fig S1), making this system ideally suited to discerning scenarios of top-down versus bottom-up co-selection. The dominant or most abundant species (Fig. 1C-D) comprises most of the biomass of these communities (median = 46%, range: (23%, 91%); N=15) (Fig. S1). Thus, we will focus on the dominants, and ask whether they co-select and are co-selected by the sub-dominant species in their communities (henceforth collectively referred to as their “cohorts”(Fig. 1C-D)). As we describe below, our results confirm that positive top-down co-selection is indeed common, but its effects are weak. In contrast, bottom-up co-selection can be very strong, and positive co-selection is far more common than negative co-selection. Our findings thus make the case that collective interactions play an important role at dictating community structure during community coalescence.

## Results and Discussion

To test the “top-down” co-selection hypothesis, we worked with eight natural microbiomes from different soil and plant environmental sources (Materials and Methods) which had been previously stabilized in serial batch-culture bioreactors for 84 generations in synthetic minimal media containing either glutamine or citrate as the only supplied source of carbon (Fig. 1A-B)^18^. We isolated the dominant species from all of these communities and identified them by Sanger-sequencing their 16S gene, which correctly matched the dominant Exact Sequence Variant (ESV)^19,20^ found through community-level 16S illumina sequencing (Fig. 1C-D and Fig. S1). In general, the dominant members of these communities remained at high frequency and were still dominant after seven additional transfers (Fig. S1), with the exception of two of the citrate communities (where the dominants went extinct and were presumably a transiently dominating species), and one of the glutamine communities (where its dominant species frequency dropped down substantially at the end of the seven extra transfers). Because studying dominant-cohort interactions is conditioned on a correct identification of the dominant, we excluded these from further analysis. For three of the citrate communities, the respective dominants had the same 16S sequence as well as a similar colony morphology; pairwise competitions between these dominants were also excluded from the analysis, as we could not easily discriminate between them.

### Top-down co-selection is weak

According to the top-down cohesiveness hypothesis formulated above, where the dominant species co-selects the rarer members of its community during coalescence (Fig. 1E; left panels), the outcome of community coalescence would be predicted by which of the two dominant species is most competitive in pairwise competition (Fig. 2A). The outcome of community coalescence may be quantified, for instance by the similarity between the coalesced and the invasive communities (as quantified by the Bray-Curtis (Fig. 2B) or Jaccard (Fig. 2C) distances), or by the relative probability of the endemic invasive species to be found in the coalesced community (Fig.2D).

**Figure 2.**
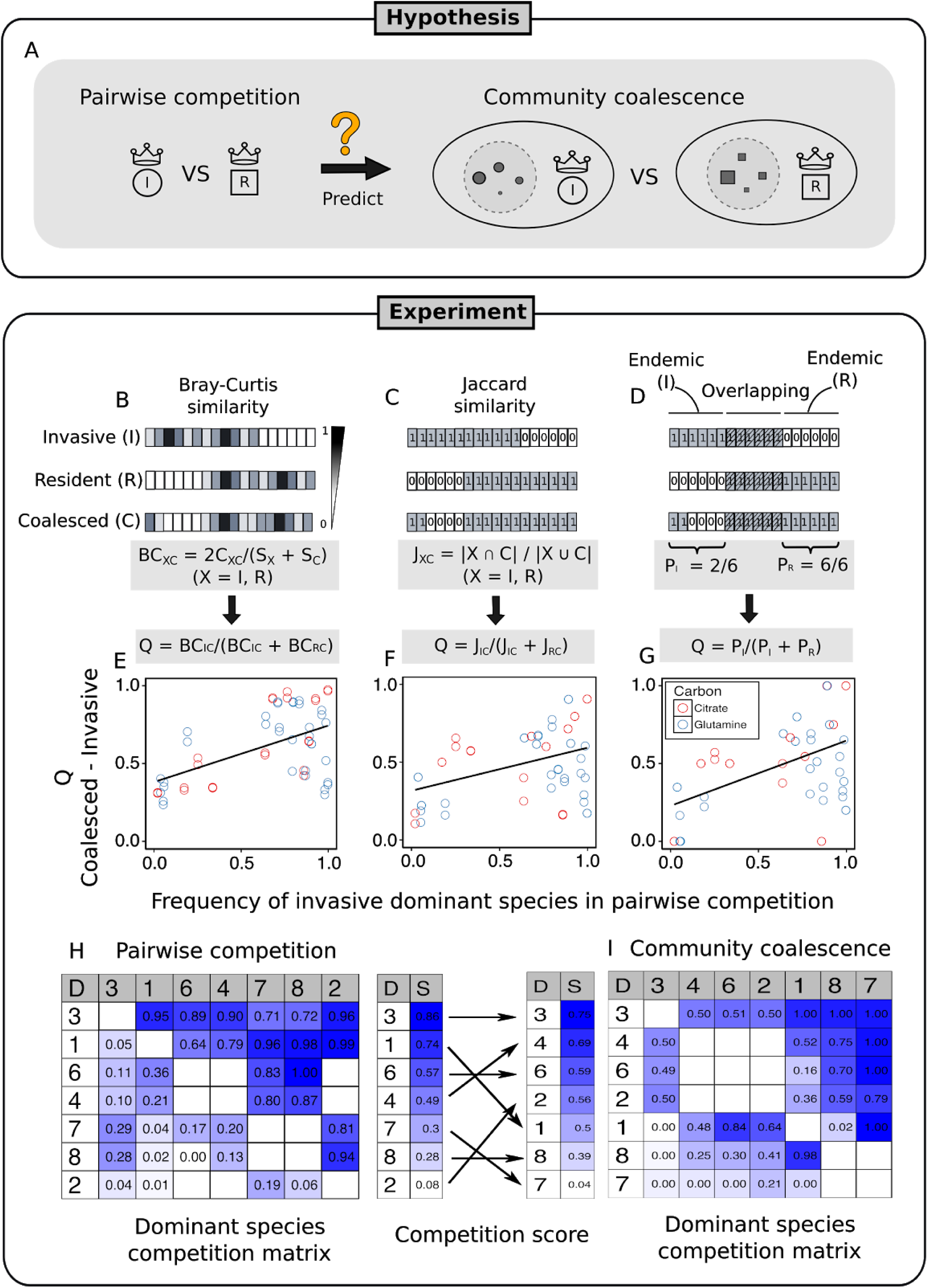
Determining the strength of top-down co-selection in our coalescence experiments. (A) In a top-down co-selection scenario, the cohort is sustained by cross-feeding from its dominant species. In such instance, pairwise competition between the dominant species in the invasive and resident communities will be predictive of the outcome of community coalescence. (B-D) This outcome is defined by the similarity between the coalesced and the invasive communities, which can be quantified by various distance metrics, e.g. the relative Bray-Curtis distance (B), relative Jaccard distance, and (C) relative survival of the endemic invasive species. (E-G) Experimental measurement of all three metrics as a function of the frequency reached by the invasive dominant in competition against the resident dominant. Each data point is calculated from four different experiments: (1) a coalescence experiment between two communities; (2) a pairwise-competition experiment between the dominants of the same two communities; (3-4) The invasive and resident communities propagated in the absence of coalescence as a control (see B-D). (H) Competition matrix for the dominant species. Numbers represent the frequency reached by the species in the row in competition against the species in the column. The average competition score for each dominant, ranked from top to bottom. (I) Dominant species competition matrix in community coalescence. Numbers represent the frequency reached by the dominant of the community in the row, relative to the frequency of the dominant species of the community in the column.

To test this hypothesis, we performed all possible pairwise competitions between dominant species in glucose and citrate environments, by mixing them 1:1 on their native media and propagating them for seven serial transfers (42 generations) (Methods). The competitive ability of a dominant species against another dominant was determined by its relative frequency after seven transfers in pairwise competition (see Methods). We also performed all possible pairwise community coalescence experiments, by mixing equal volumes of all communities, and propagating the coalesced communities for an extra seven transfers (Fig. 1F). The frequencies of all species in both community-community and dominant-dominant competitions were determined through 16S illumina sequencing (Methods). In total, we include 56 community coalescence and 56 dominant pairwise competitions in our analysis. Compiling the results from all of these experiments, we found that the pairwise competitive ability of an invasive dominant is only weakly predictive (though statistically significant) of the performance of its community during community coalescence, as quantified by the relative Bray-Curtis similarity between the coalesced and the invasive communities (adjusted R^2^ = 0.26, P<0.001; Fig. 2E), and, although the effect is even weaker, also by the relative Jaccard similarity between the coalesced and the invasive communities (adjusted R^2^ = 0.15, P = 0.0017; Fig. 2F). These two metrics include the presence of the dominant species itself. To better disentangle the effect that the dominants have on the other members of their communities, we excluded the dominant species from the compositional data, and found that our results still hold (Bray-Curtis R^2^ = 0.17, P<0.001; Jaccard R^2^ = 0.11; P=0.007; Fig. S2). To further characterize the extent to which dominant competition governs community coalescence, we also determined the relative invasion success of the endemic invasive species (i.e. those that are present in the invasive but not the resident community). We found that the relative frequency of a dominant against another in head-to-head pairwise competition is only weakly predictive of the relative success of its endemic community members during community coalescence (linear regression, adjusted R^2^ = 0.23, P< 0.001; Fig. 2G).

The head-to-head pairwise competitive interactions amongst the dominants were very hierarchical (Fig. 2G; Fig. S3), exhibiting perfect transitivity (See Materials and Methods). However, when we estimated the dominant-dominant competition coefficients in the presence of their cohorts (i.e. by calculating the relative frequency of one dominant against the other after community coalescence (See Methods)), we found that the competitive hierarchy changed substantially, and transitivity was violated as a result of the presence of the cohorts (Fig. 2G-H). Likewise, the relative frequency of a dominant against another in head-to-head pairwise competition is only weakly (linear regression R^2^ = 0.28; P=0.002; Fig. S4) predictive of its relative frequency against that same other dominant when the cohorts are present too (e.g. after coalescence).

Together, these results suggest that, although some level of top-down co-selection by the dominants towards their cohorts is consistent with our experiments, the cohorts are not just passively responding to their dominants, and their fates are not strictly determined by dominant-dominant competition. Rather, our evidence suggests that the cohorts may be playing an active role in community coalescence, significantly affecting the fate of their dominants. This finding led us to investigate the potential role of bottom-up ecological co-selection (Fig. 1E, right panels), in which the dominants may be co-selected for or against by their cohorts during community coalescence.

### Bottom-up co-selection during microbial community coalescence

The effect of bottom-up co-selection of the dominant by the cohort can be visualized by comparing the invasion success of the dominant from community (I) invading community (R) alone with its success when it invades accompanied by its cohort (Fig. 3A). We delineate two limiting scenarios. First, strong positive bottom-up co-selection (recruitment) would be represented by the green region along the vertical axis in Fig. 3A. These are hypothetical cases where a dominant cannot invade a resident community alone, but it is able to and may even reach high frequencies when co-invading with its cohort. In the other extreme, represented by the red shaded region along the horizontal axis (Fig. 3A), co-selection occurs in a negative direction: an invasive dominant that alone is able to successfully invade a resident community but fails to do so when it invades accompanied by its resident cohort, hampered by it.

**Figure 3.**
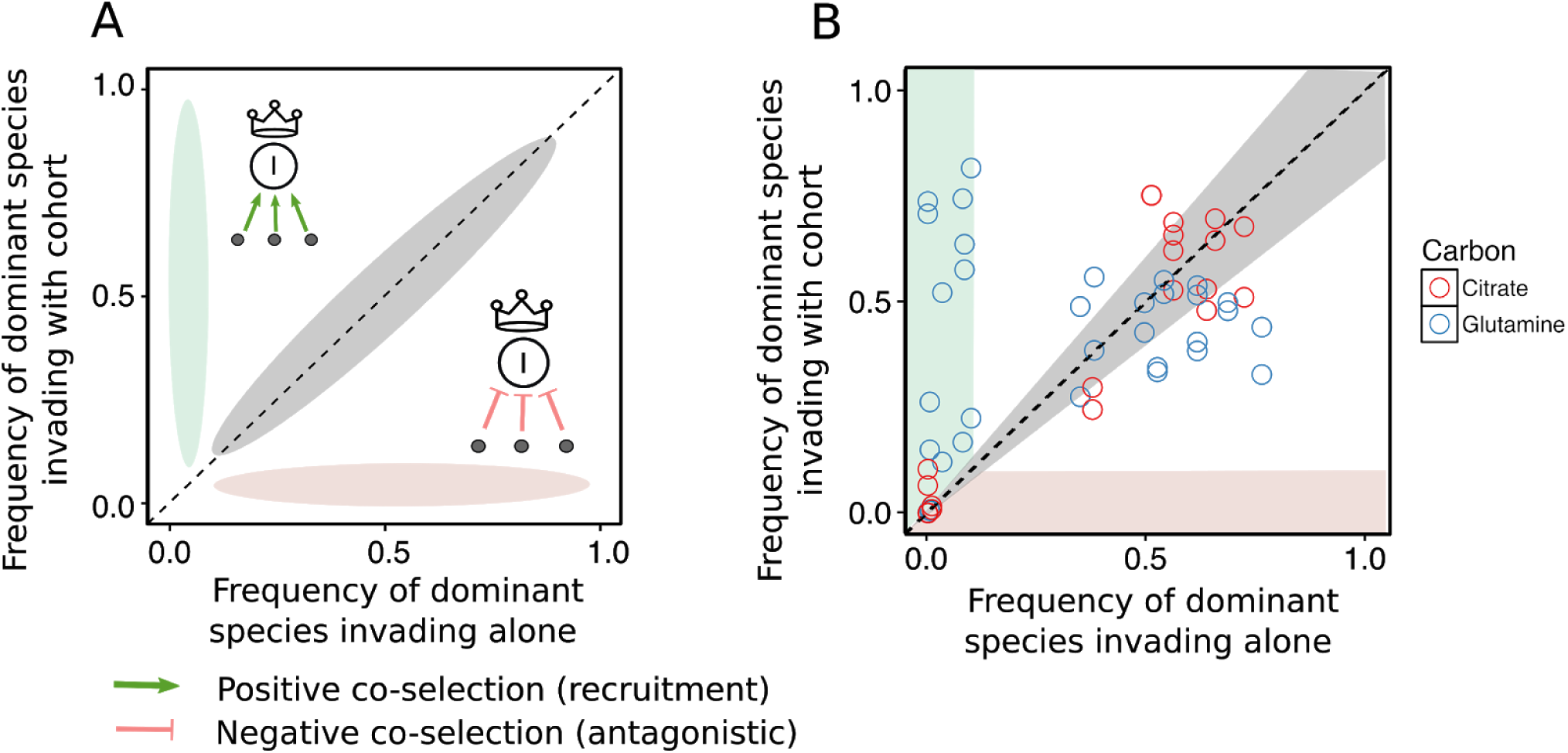
Evidence of bottom-up co-selection in microbial community coalescence. (A) Three potential scenarios of bottom-up co-selection. Green and red shaded regions represent limit cases of positive (recruitment), or negative (antagonistic) bottom-up co-selection. Gray region along the diagonal would represent dominant species that reach the same frequency when they invade the resident community alone as they do when they invade it in the company of their cohort. (B) Experimental comparison of the frequency reached by invasive dominants against resident communities when invading alone, or as a part of their native community shows that positive bottom-up co-selection is common, but antagonistic co-selection is rare in our experiments.

To populate an experimental plot akin to that in Fig. 3A, we performed a new round of invasion experiments, where each dominant invaded every community in isolation (See Methods section for details). The dominant-invaded communities were then propagated for seven serial transfers, and the final community compositions were determined by Illumina 16S sequencing. The regions of strong positive and negative bottom-up co-selection are defined by the shaded green and red areas in Plot 3B, which correspond to frequencies lower than 0.1 when invading alone (positive co-selection; green), or with the cohort (negative co-selection; red). A significant number of dominants (13/24 or 54%) that were not able to invade a resident community alone (or reached a frequency of at most 0.1) were able to do so and even reach very high frequencies when invading with their cohorts, indicating that positive bottom-up co-selection is indeed at play and common in our experiments. In contrast, negative co-selection is rare (Fig. 3B; green shaded area), as no instances were observed. A substantial fraction of dominants (17/56 or 30%) lay within 20% of the identity line (gray region), suggesting that whether the invasive cohort is present or not during invasion does not significantly impact the invasive dominant’s invasive success.

## Discussion

Understanding the responses of microbial communities to invasions, and the processes leading to them is an essential but poorly understood question in microbial ecology ^10^. Theory has long suggested that communities may exhibit “cohesiveness” in the face of invasions^3,4,14,15^, and that members of the same community can co-recruit (or co-select) one another during community-community invasions. Our results provide direct experimental evidence of ecological co-selection in a large number of microbial community coalescence experiments. Our results highlight the critical role played by the rarer, sub-dominant species to generating community cohesiveness during community coalescence.

A recent study has also reported indirect evidence of ecological co-selection in a different experimental system (methanogenic communities in anaerobic single-batch digesters inoculated with industrial or agricultural waste), where facilitation is also believed to be strong ^12^. Given the convergence of results between different experiments with different communities and in very different environments, and the confluence with theoretical predictions ^3^, we conclude that collective interactions arising from metabolic feedbacks between microbial growth and the environment should be generically expected to produce ecological co-selection of members of the same community during community coalescence. The experimental system introduced here can be easily expanded, and hundreds or even thousands of community coalescence experiments can be carried out in parallel. It thus represents a promising tool to explore the generic properties of microbial community coalescence in high throughput, and to test quantitative theories about its role in microbiome assembly.

## Methods

### Stabilization of environmental communities in simple synthetic environments

Communities were stabilized ex situ as described elsewhere ^18^. Briefly, environmental samples (soil, leaves) within one meter radius in eight different geographical locations were collected with sterile tweezers or spatulas into 50 mL sterile tubes (Fig. 1A). One gram of each sample was allowed to sit at room temperature in 10 mL of phosphate buffered saline (1xPBS) containing 200µg/mL cycloheximide to suppress eukaryotic growth. After 48h, samples were mixed 1:1 with 80% glycerol and kept frozen at −80°C. Starting microbial communities were prepared by scrapping the frozen stocks into 200 µL of 1xPBS and adding a volume of 4µL to 500µL of synthetic minimal media (1xM9) supplemented with 200µg/mL cycloheximide and 0.07 C-mol/L glutamine or sodium citrate as carbon sources in 96 deep-well plates (1.2 mL; VWR). Cultures were then incubated still at 30°C to allow re-growth. After 48h, samples were fully homogenized and biomass increase was followed by measuring the optical density (620nm) of 100µL of the cultures in a Multiskan FC plate reader (Thermo Scientific). Microbial communities were stabilized ^18^ by repeating every 48h the passage of 4 µL of the cultures into 500 µL of fresh media (1xM9 and carbon source) for a total of 12 transfers at a dilution factor of 100X, roughly equivalent to 80 generations per culture (Fig. 1B). Cycloheximide was eliminated from the media after the two first transfers.

### Determination of community composition by 16S sequencing

The protocol was identical to that followed elsewhere ^18^. Community samples were collected by spinning down at 3,500 rpm for 25 min in a bench-top centrifuge at room temperature; cell pellets were stored at −80 °C before processing. To maximize Gram-positive bacteria cell wall lysis, the cell pellets were re-suspended and incubated at 37 °C for 30 min in enzymatic lysis buffer (20 mM Tris-HCl, 2mM sodium EDTA, 1.2% Triton X-100) and 20 mg/mL of lysozyme from chicken egg white (Sigma-Aldrich). After cell lysis, the DNA extraction and purification was performed using the DNeasy 96 protocol for animal tissues (Qiagen). The clean DNA in 100 µL elution buffer of 10 mM Tris-HCl, 0.5 mM EDTA at pH 9.0 was quantified using Quan-iT PicoGreen dsDNA Assay Kit (Molecular Probes, Inc.) and normalized to 5 ng/µL in nuclease-free water (Qiagen) for subsequent 16S rRNA illumina sequencing. 16S rRNA amplicon library preparation was performed following a dual-index paired-end approach ^21^. Briefly, PCR amplicon libraries of V4 regions of the 16S rRNA were prepared using dual-index primers (F515/R805), then pooled and sequenced using the Illumina MiSeq chemistry and platform. Each sample went through a 30-cycle PCR in duplicate of 20 µL reaction volumes using 5 ng of DNA each, dual index primers, and AccuPrime Pfx SuperMix (Invitrogen). The thermocycling procedure includes a 2-min initial denaturation step at 95 °C, and 30 cycles of the following PCR scheme: (a) 20-second denaturation at 95 °C, (b) 15-second annealing at 55 °C, and (c) 5-min extension at 72 °C. The duplicate PCR products of each sample were pooled, purified, and normalized using SequalPrep PCR cleanup and normalization kit (Invitrogen). Barcoded amplicon libraries were then pooled and sequenced using Illumina Miseq v2 reagent kit, which generated 2X250 base pair paired-end reads at the Yale Center for Genome Analysis (YCGA).

The sequencing reads were demultiplexed on QIIME 1.9.0 ^22^. The barcodes, indexes, and primers were removed from raw reads, producing FASTQ files with both the forward and reverse reads for each sample, ready for DADA2 analysis ^19^. DADA2 v. 1.1.6 was used to infer unique biological exact sequence variants (ESVs) for each sample and naïve Bayes was used to assign taxonomy using the SILVA v. 123 database ^23,24^.

### Isolation of dominants from all communities

For each community, the most abundant colony morphotype at the end of the ninth transfer was selected, resuspended in 100µL 1xPBS and serially diluted (1:10). Next, 20µL of the cells diluted to 10^-6^ were plated in the corresponding synthetic minimal media and allowed to regrow at 30°C for 48h. Dominants were then inoculated into 500µL of fresh media and incubated still at 30°C for 48h. After this period, the communities stabilized for eleven transfers and the isolated dominants were ready for the competition experiments (Fig 1F) at the onset of the twelfth transfer.

### Dominant-dominant and community-community competitions

All possible pairwise dominant-dominant and community-community competition experiments were made by mixing equal volumes (4 µL) of each of the eight communities or eight dominants at the onset of the twelfth transfer. Competitions were set up on their native media, *i.e.* in 500 µL of 1xM9 supplemented with either 0.07 C-mol/L of glutamine or citrate in 96 deep-well plates, and were incubated at 30°C for 48h. Pairwise competitions were further propagated for seven serial transfers (roughly 42 generations; Fig 1F) by transferring 8µL of each culture to fresh media (500 µL).

### Determining average competitive scores and hierarchy scores

The average competitive scores and hierarchy scores of the dominant-dominant (d-d) and community-community (c-c) competitions were computed according to L. M. Higgins et al ^25^. The average competitive score 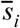 of each dominant species *i* was defined as its mean frequency *f* _*ij*_ after pairwise dominant-dominant competitions with each of the *n* − 1 dominant species or after invading each of the *n* − 1 community *j* during dominant-community competitions or coalescence of community *i* with each of the *n* − 1 communities:

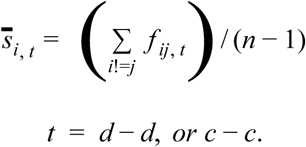

The competitiveness of each dominant strain during the d-d or c-c competitions were ranked based on the average competitive scores under each type of competition.

### Metrics of community distance

(Jaccard and Bray Curtis formulae)

Beta-diversity indexes between the invasive and the coalesced communities or the resident and the coalesced communities were performed using Bray–Curtis or Jaccard similarity metrics, as implemented in R package Vegan ^26^). The fractions of the endemic cohort from the invasive communities that persisted in the coalesced communities were also used to compute community distances.

## Acknowledgements

The authors wish to thank Joshua Goldford, Pankaj Mehta, Wenping Cui, Robert Marshland, and all members of the Sanchez laboratory for many helpful discussions. We also wish to express our gratitude to the Goodman laboratory at Yale for technical help during the early stages of this project. The funding for this work partly results from a Scialog Program sponsored jointly by Research Corporation for Science Advancement and the Gordon and Betty Moore Foundation through grants to Yale University by the Research Corporation and by the Simons Foundation.

## Supplementary Figures

**Figure S1.**
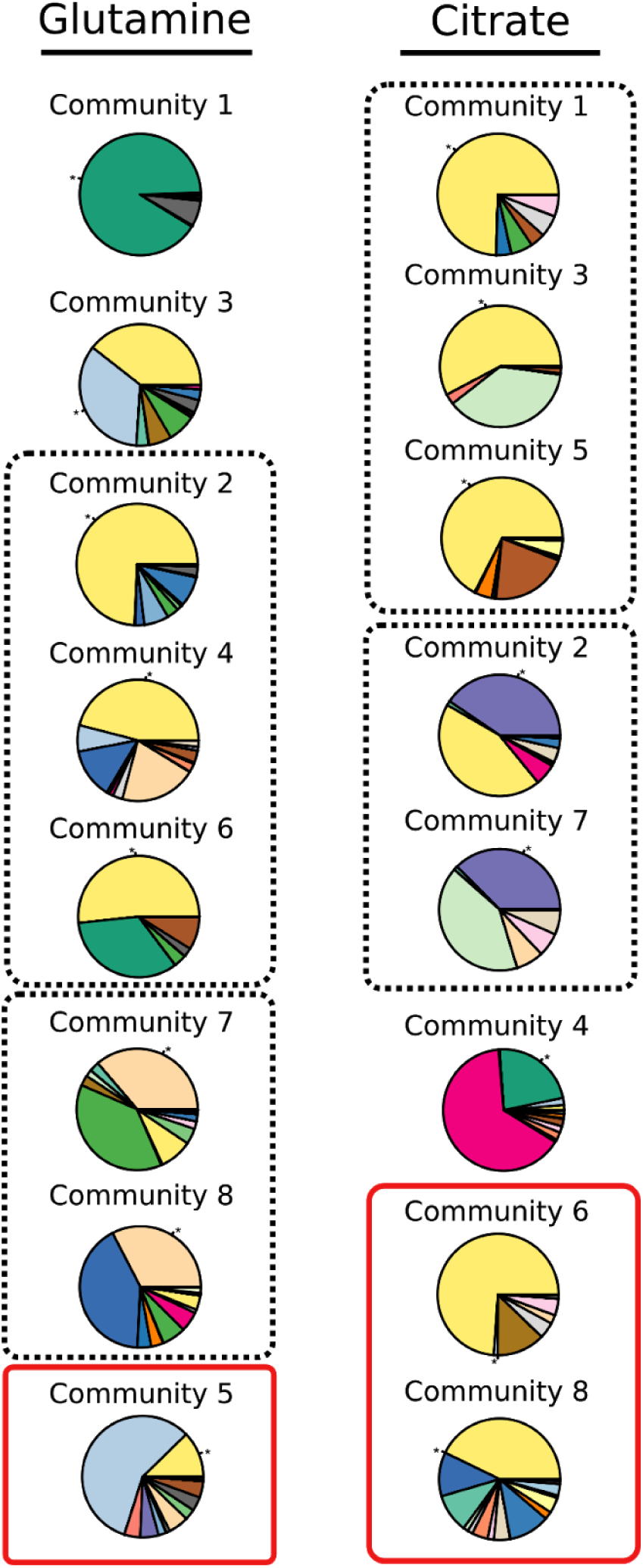
Invasive or resident community compositions without coalescence after seven transfers. The dominant members of most communities remained at high frequency and were still dominant after seven additional transfers. Each color slide in the pie chart indicates one exact sequence variant (ESV). The dominant species ESV isolated by plating is labeled with asterisk. The communities clustered by the dashed boxes shared the same ESV as dominant species. The communities highlighted by red boxes had the dominant species isolated by plating not matching the sequencing results, therefore, were eliminated from analyses.

**Figure S2.**
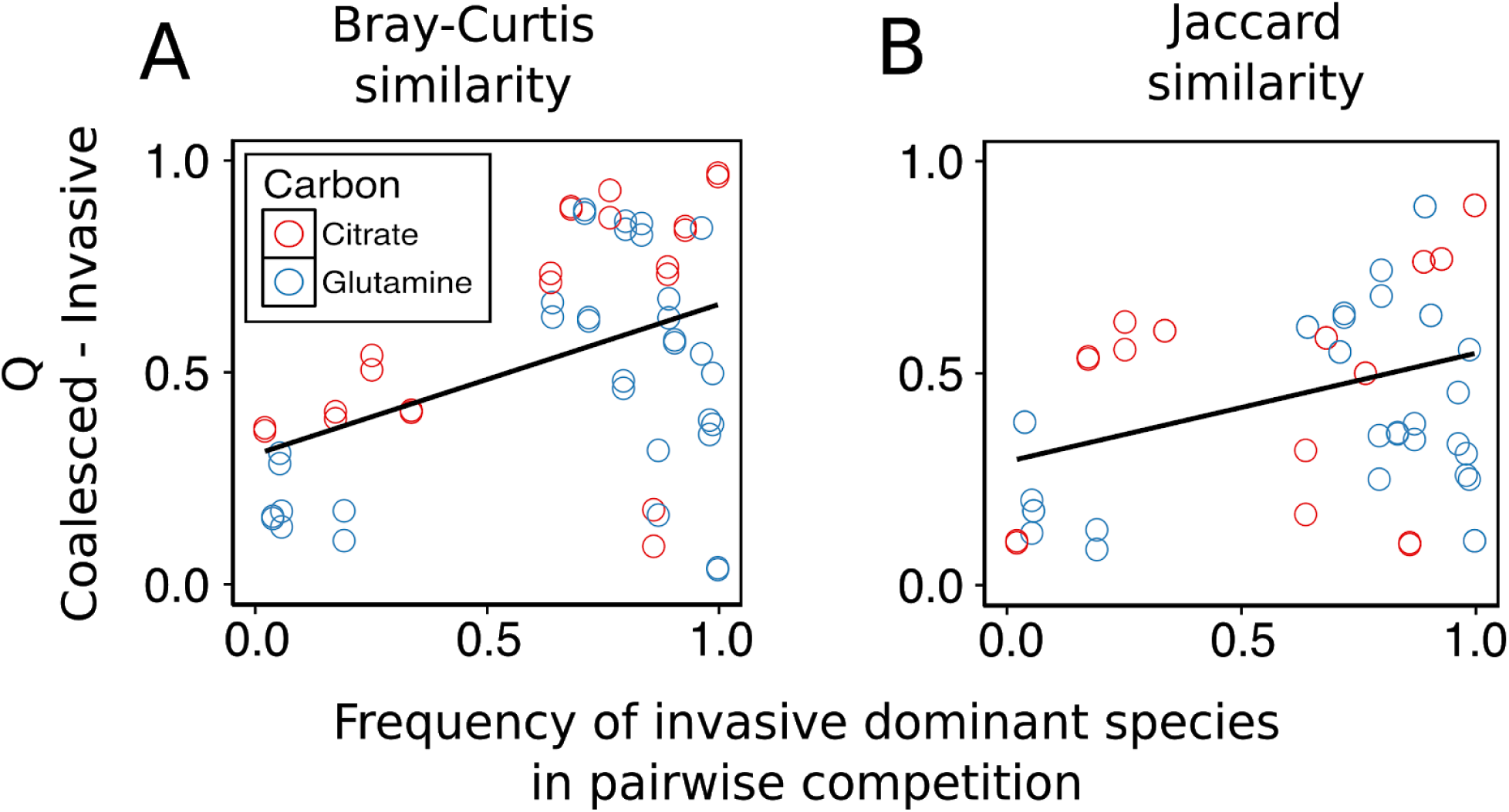
Limited effect of the dominants have on the coalescence results. We excluded the dominant species from the compositional data, and repeated our analysis in Fig. 2. A significant (though weaker) effect is found both for Jaccard and Bray-Curtis, indicating that the dominants apply top-down co-selection (recruitment) towards their cohort.

**Figure S3.**
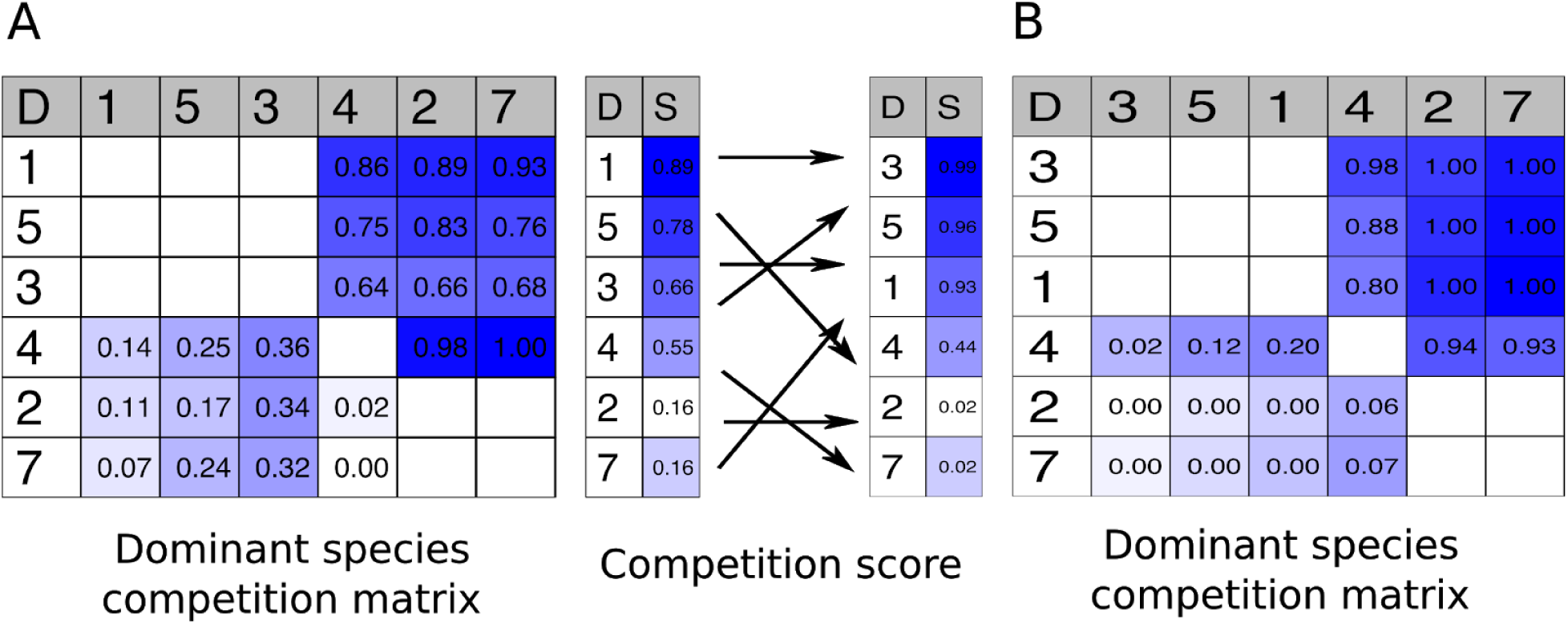
Dominant species competition matrix for the communities assembled in citrate.

**Figure S4.**
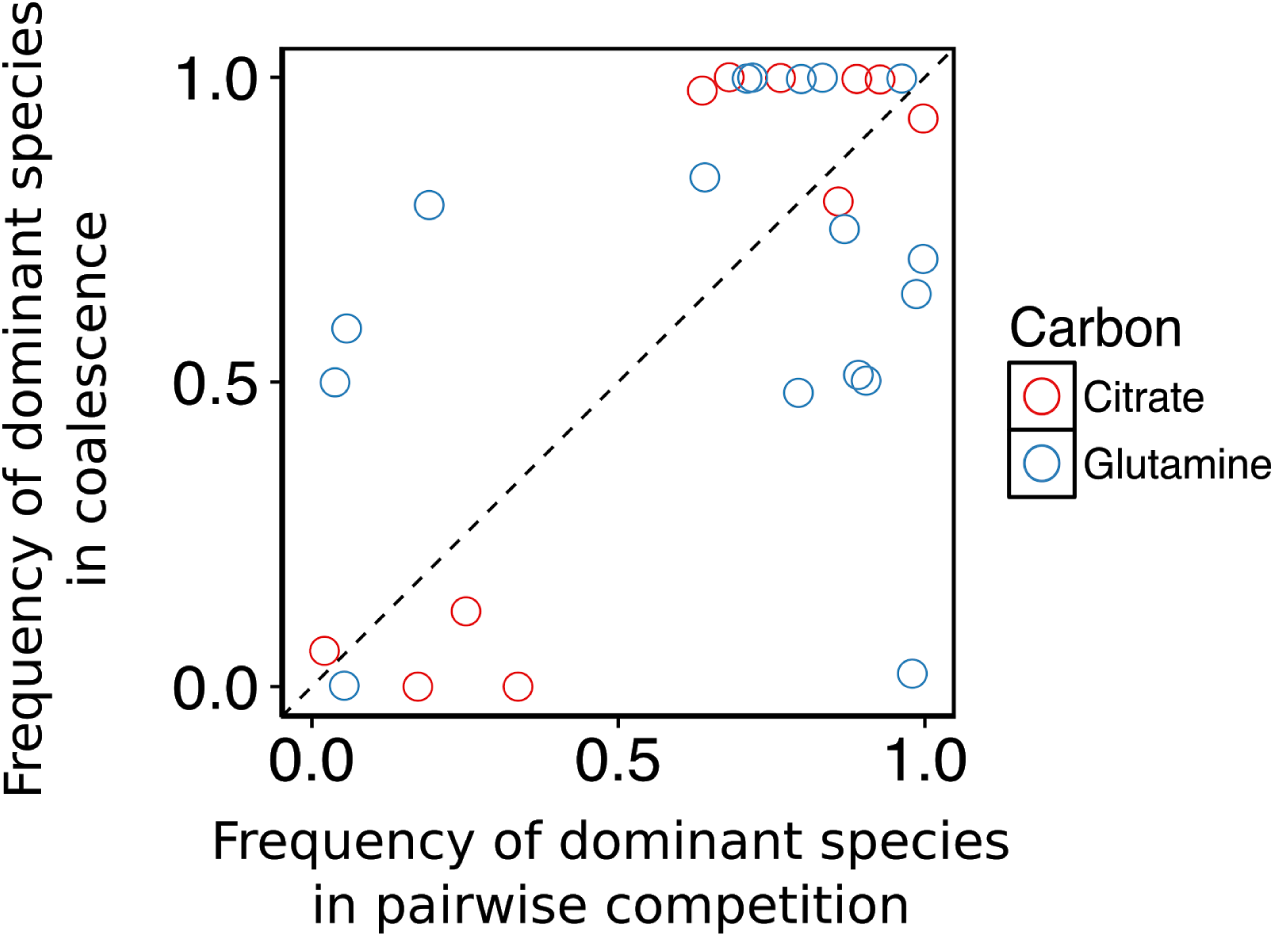
We plot the relative frequency of a dominant species against another dominant either in direct head-to-head or during community coalescence. We find that the former is a poor predictor of the latter (linear regression R^2^ = 0.28; P=0.002; Fig. S4), suggesting that the cohort is not simply cross-fed by their dominant taxon, but can itself influence the competitive ability of their dominant. Each data point is an average of two replicates.

